# Gut commensal bacteria enhance pathogenesis of a tumorigenic murine retrovirus

**DOI:** 10.1101/2022.02.02.478820

**Authors:** Jessica Spring, Sophie Lara, Aly A. Khan, Kelly O’Grady, Jessica Wilks, Sandeep Gurbuxani, Steven Erickson, Michael Fischbach, Amy Jacobson, Alexander Chervonsky, Tatyana Golovkina

## Abstract

The influence of the microbiota on viral transmission and replication is well appreciated. However, its impact on retroviral pathogenesis outside of transmission/replication control remained unknown. Using Murine Leukemia Virus (MuLV), we found that some commensal bacteria promoted the development of leukemia induced by this retrovirus. The promotion of leukemia development by commensals was due to suppression of the adaptive immune response through upregulation of several negative regulators of immunity. These negative regulators included Serpinb9b and Rnf128, which are associated with a poor prognosis of some spontaneous human cancers. Upregulation of Serpinb9b was mediated by sensing of bacteria by NOD1/NOD2/RIPK2 pathway. This work describes a novel mechanism by which the microbiota enhances tumorigenesis within gut-distant organs and points at potential new targets for cancer therapy.

## Introduction

The commensal microbiota plays a critical role in maintaining host health by providing nutrition, creating a hostile environment for incoming bacterial pathogens, and modulating maturation of secondary lymphoid organs (Kim et al., 2017; Littman and Pamer, 2011). Importantly, microbiota is among the factors influencing development of tumors. The gut microbiota has been implicated in progression of cancers of the gut and associated organs. That effect is achieved by induction of inflammation through pattern recognition receptors and metabolites that lead to induction of inflammatory cytokines and chemokines, or through production of DNA-damaging reactive oxygen and nitrogen species (Arthur et al., 2012; Hope et al., 2005; Rakoff-Nahoum and Medzhitov, 2007; Reddy et al., 1975; Yoshimoto et al., 2013). The gut microbiota can also be tumor suppressive (Donohoe et al., 2014) or enhance the effect of anti-cancer immunotherapy by modifying dendritic cell maturation and promoting tumor specific T cell responses (Sivan et al., 2015; Vetizou et al., 2015).

Some oncogenic retroviruses take advantage of the microbiota for their spread and replication. Previously, we demonstrated that transmission of Mouse Mammary Tumor Virus (MMTV) that causes mammary carcinomas in susceptible mice via insertional mutagenesis depends on the gut commensal bacteria (Kane et al., 2011). Orally transmitted MMTV binds bacterial lipopolysaccharide (LPS) through LPS binding receptors incorporated into the viral membrane during budding (Wilks et al., 2015). LPS attached to the viral membrane triggers TLR4 to induce inhibitory cytokine interleukin-10 (IL-10), thus generating a status of immunological tolerance to the virus (Kane *et al*., 2011). Therefore, the microbiota enables successful virus replication and thus, subsequent tumorigenesis by suppressing the anti-viral immune response.

In this study, we sought to determine whether the commensal microbiota influences replication and/or pathology induced by a retrovirus capable of transmitting through the blood – Murine Leukemia Virus (MuLV) (Buffett et al., 1969; Duggan et al., 2006). This virus induces leukemias in susceptible mice (Friend, 1957; Rauscher, 1962). To elucidate the role of the microbiota on MuLV infection and associated pathology, Rauscher like-MuLV (RL-MuLV), which causes erythroid leukemia in mice from susceptible backgrounds (Hook et al., 2002) was used. Here, we established that while the gut microbiota had no effect on replication and transmission of the virus, it significantly enhanced virally-induced leukemia development. The tumor progression was aided by select gut commensal bacteria, which upregulated signaling pathways counteracting the immune response capable of restricting leukemia development.

## Methods

### Mice

Mice utilized in this study were bred and maintained at the animal facility of The University of Chicago. BALB/cJ mice were purchased from The Jackson Laboratory (TJL). C.129S7(B6)-*Rag1^tm1Mom^*/J, C.C3-*Tlr4^Lps-d^*/J, C.129(B6)-*Tlr2^tm1Kir^*/J, and B6.129S1-*Ripk2^tm1Flv^*/J mice were purchased from TJL and backcrossed to BALB/cJ mice for 10 generations to create RAG1^-/-^, TLR4^-/-^, TLR2^-/-^, and RIPK2^-/-^ mice, respectively, on the BALB/cJ background. TLR2^-/-^ and TLR4^-/-^ mice on the BALB/cJ background were intercrossed to produce double deficient mice. CByJ.129S2(B6)-IL6^tm1Kopf^/J, purchased from TJL were crossed to BALB/cJ mice and the resulting F1 mice were intercrossed to generated -/- and +/+ littermate mice used in the studies.

Serpinb9b^-/-^, Rnf128^-/-^, and VSig4^-/-^ BALB/cJ mice were generated using the CRISPR/Cas9 approach. To generate VSig4^-/-^ mice, two guide RNAs targeting exon 1 (5’AGAAGTAGCTTCAAATAGGA3’ and 5’TGAGCACTATTAGGTGGCCC3’) were used. Five distinct BALB/cJ VSig4^-/-^ lines were produced. Primers flanking exon 1 (5’CCAGAACAAATGGTTTCCCTAG3’ and 5’TTGGAGTCCTCT-GACATCCC3’) were used to genotype the founder mice and subsequent offspring. Equal numbers of mice from each line were used in all experiments.

To generate Serpinb9b^-/-^ mice, two guide RNAs were used to target exon 2 (5’TATGGTCCTCCTGGGTGCAA3’ and 5’TTCCACCCTAGTTGTTACTC3’). Genotyping was performed using two primers flanking exon 2 (5’TTTCCCTCCAGACTCTGCA3’ and 5’GAGAAGCATGAGGCCAAGAC3’). Two lines of Serpinb9b^-/-^ mice were produced. Equal numbers of mice from both lines were used in all experiments. For the generation of BALB/cJ Rnf128^-/-^ mice, one guide RNA was used to target exon 4 (5’AGGATGCGCACCAAATCATT3’). One line of Rnf128^-/-^ mice was developed. Founder VSig4^-/-^, Serpinb9b^-/-^, and Rnf128^-/-^ mice were crossed to BALB/cJ mice for at least two generations. Heterozygous mice were then intercrossed to produce homozygous knockout (KO) animals used in subsequent studies.

Males and females were used at ~50:50 ratio in all experiments. The studies outlined here have been reviewed and approved by the Animal Care and Use Committee at The University of Chicago.

### Antibiotic treatment

BALB/cJ females injected intraperitoneally (i.p.) with 1 x 10^3^ plaque forming units (pfus) of RL-MuLV were bred with BALB/cJ males to produce infected G1 animals. The G1 mice were weaned onto water containing the following antibiotic mixture: kanamycin (4 mg/ml), gentamicin (3.5 mg/ml), colistin (8500 U/ml), metronidazole (2.15 mg/ml), and vancomycin (4.5 mg/ml) for the duration of the experiment. When the mice became symptomatic (anemic, lethargic, enlarged abdomen), they were sacrificed and assessed for leukemia development. All animals were sacrificed by 5 months of age. The efficacy of the antibiotic-treatment protocol was evaluated by periodical bacteriologic examination of feces. Fresh fecal samples from antibiotic-treated and control mice were vortexed in TSA broth (0.1g in 1ml) and plated on BBL plates. Plates were incubated for 48h at 37°C under both aerobic and anaerobic conditions. Anaerobic growth conditions were created using a COY anaerobic chamber (Grass Lake, MI). There were no detectable colony forming units (CFUs) on either aerobic or anaerobic plates from the antibiotic treated animals. The plates from the untreated animals had 10^7^ CFUs and 8 x 10^5^ CFUs of anerobic and aerotolerant bacteria per 1g of feces, respectively.

### Monitoring sterility in GF isolators

BALB/cJ and BALB/cJ RAG1^-/-^ mice were re-derived as germ-free (GF) at Taconic and housed in sterile isolators at the gnotobiotic facility at the University of Chicago. Sterility of GF isolators was assessed as previously described (Wilks et al., 2014). Briefly, fecal pellets were collected from isolators weekly and quickly frozen. DNA was extracted using a bead-beating/phenol-chloroform extraction protocol. A set of primers that broadly hybridize to bacterial 16S rRNA gene sequences were used to amplify isolated DNA. In addition, microbiological cultures were set up with GF fecal pellets, positive control SPF fecal pellets, sterile saline (sham), and sterile culture medium (negative control). BHI, Nutrient, and Sabbaroud Broth tubes were inoculated with samples and incubated at 37°C and 42°C aerobically and anaerobically. Cultures were followed for five days until cultures were deemed negative.

### Colonization of GF mice

GF BALB/cJ mice were colonized with the ASF consortium (Sarma-Rupavtarm et al., 2004) via oral gavage of suspended cecal contents from donor mice. All but one species (ASF360, *Lactobacillus acidophilus*) were found in the ASF-colonized mice. *Lactobacillus murinus* (*L. murinus*, ASF361) and *Parabacteroides goldsteinii* (*P. goldsteinii*, ASF519) were isolated from the ASF consortium. *L. murinus* was isolated by plating suspended fecal matter from ASF-colonized mice onto de Man, Rogosa and Sharpe agar (MRS) (Thermo Fisher Scientific), a selective media for *Lactobacilli*. Growth of *L. murinus* was confirmed by PCR. *P. goldsteinii* was isolated by plating fecal matter from ASF colonized mice re-suspended in PBS onto brain heart infusion supplemented (BHIS) media, supplemented with hemin and vitamin K1, and grown in an anaerobic chamber. Growth of *P. goldsteinii* was confirmed by PCR. *Bacteroides thetaiotaomicron* (*B. theta*, VPI-5482 Δtdk) was grown on tryptone yeast extract glucose (TYG) media in an anaerobic container. *L. murinus, P. goldsteinii*, and *B. theta* were introduced to GF BALB/cJ mice by oral gavage of 200μl of overnight liquid culture grown from single colonies. Successful colonization was confirmed via PCR using primers specific for each individual bacterium. Female offspring of colonized mice were infected with filter-sterilized virus (G0 mice), bred to produce G1 mice which were monitored for leukemia

### Virus isolation, infection, and leukemia monitoring

Rauscher-like Murine Leukemia Virus (RL-MuLV) which consists of N, B tropic ecotropic and mink cell focus forming (MCF) virus (Hook *et al*., 2002) was isolated from tissue culture supernatant of infected SC-1 cells (ATCC CRL-1404). The virus was tested for hepatitis virus, mouse thymic virus, mouse parvovirus, pneumonia virus of the mouse, polyomavirus, mammalian orthoreovirus serotype 3, enzootic diarrhea virus of infant mice, Sendai virus, *Mycoplasma (M) pulmonis*, *M. arginine*, *M. arthritidis*, *M. bovis*, *M. cloacale*, *M. falconis*, *M. faucium*, *M. fermentans*, *M. genitalium*, *M. hominis*, *M. hyorhinis*, *M. hyosynoviae*, *M. opalescens*, *M. orale*, *M. pirum*, *M. pneumonia*, *M. salivarium*, *M. synoviae*, *Acholeplasma laidlawii*, *Ureaplasma urealyticum*, cilium-associated respiratory bacillus, ectromelia virus, encephalitozoon cuniculi, Theiler’s murine encephalomyelitis virus, Hanta virus (Korean hemorrhagic virus), lymphocytic choriomeningitis virus, lactate dehydrogenase enzyme, minute virus of the mouse, mouse adenovirus, and mouse cytomegalovirus and was found to be negative for all the pathogens.

Ecotropic virus titers in the RL-MuLV mixture were determined by an XC infectious plaque assay as described in (Rowe et al., 1970). 1 x 10^3^ pfus of the virus isolated from supernatants of infected SC-1 cells were injected i.p. into 6-8 week-old BALB/cJ mice (G0 mice). Mice were bred to produce G1 animals which were used as a source of the virus for all experiments. The virus was not passaged beyond G1 *in vivo*. To isolate the virus, splenic homogenates of non-leukemic 2-3-month-old G1 mice were centrifuged at 2,000 rpm for 15 min at 4°C. The supernatant was layered onto a 30% sucrose cushion and spun down at 31,000 rpm using a TW55.1 bucket rotor for 1 hr at 4°C. The pelleted fraction was resuspended in PBS, which was spun at 10,000 rpm at 4°C to remove insoluble material. The resulting supernatant was aliquoted, titered and stored at −80°C.

Splenic derived RL-MuLV was diluted in sterile PBS and filtered through a sterile 0.22μm membrane in a laminar flow hood. 1 × 10^3^ pfus were injected i.p. into GF and SPF BALB/cJ females, which were used as controls in all experiments with gnotobiotic mice. For genetically altered mice, -/-, +/+, and in some instance +/- female littermates were injected i.p. with 1 × 10^3^ pfus. In addition, non-littermate wild-type (WT) BALB/cJ mice bred in the colony were also infected and served as non-littermate controls. Infected females were put in mating to generate G1 animals. G1 mice were aged and monitored for leukemia development which included signs of lethargy, hunched back, ruffled/unkempt fur, distended abdomen, and anemia. Symptomatic mice were sacrificed. All mice were sacrificed by 5 months, at the conclusion of the experiment. Survival curves indicate symptomatic mice that were closed during 150 days. The survival curve for SPF mice is a cumulative curve generated over the course of the entire studies using BALB/cJ mice. +/- and +/+ littermates were used for each of the experimental groups of genetically modified mice and their survival curves were compared with the cumulative curve for SPF mice to ensure that the leukemia development rate is the same (Figures S3E-F). Comparison between leukemia score and spleen weight indicate that spleen weights ≥ 0.35g have a leukemia score of 2 or higher (Figure S1D), which was used for some wild-type BALB/cJ SPF mice as a proxy for denoting leukemia. Leukemia or lack of it for all gnotobiotic, genetically modified and control SPF mice was also confirmed histologically using splenic sections.

### Histology

Mouse spleens were fixed in Telly’s fixative, sectioned, and stained with hematoxylin and eosin. Leukemia was scored using the following criteria: maintenance of splenic architecture, level of cellular mitotic activity and degree of cell maturation (Figure S1C).

### Plaque and infectious center assays

Blood from RL-MuLV infected BALB/cJ mice was collected in tubes containing heparin, which was then spun at 10,000 rpm for 10 min at 4°C. The plasma was serially diluted in PBS, and was subjected to the XC plaque assay (Rowe *et al*., 1970).

To assess the frequency of infected cells, single cell suspensions (with red blood cells) from RL-MuLV infected mouse spleens were serially diluted in Clicks medium (Irvine scientific) and were subjected to the infectious center assay (Rowe *et al*., 1970).

### Fluorescence-activated cell soring (FACS) analysis

For analysis of cells expressing hematopoietic stem cell (HSC) marker (Sca-1), single cell suspensions from mouse spleens were stained with fluorescein isothiocyanate (FITC)-conjugated anti-Ly-6A/E (Sca-1) mAb (Biolegend; San Diego, CA). To analyze erythrocyte maturation, splenocyte suspensions were stained with FITC-conjugated anti-CD71 (transferrin receptor) mAb (Biolegend; San Diego, CA) and phycoerythrin (PE)-conjugated anti-Ter119 (erythroid cells). Stages of erythrocyte maturation were determined as described in (Zhang et al., 2003). Regions R1 to R5 were defined by the following pattern: CD71^med^TER119^low^, CD71^high^TER119^low^, CD71^high^TER119^high^, CD71^med^TER119^high^, and CD71^low^TER119^high^, respectively. Dead cells (propidium iodide–positive) were excluded from the analysis. For analysis of VSig4 deletion in VSig4-deficient mice, peritoneal macrophages were stained with FITC-conjugated anti-F4/80 mab (Biolegend; San Diego, CA) and allophycocyanin (APC)-conjugated anti-VSig4 mAb (eBioscience; San Diego, CA).

### Hematopoietic stem cells (HSC) quantification in GF and SPF mice

Bone marrow was isolated from 3 SPF and 3 GF mice. The red blood cells were lysed, and 2 x 10^7^ cells were stained with the following monoclonal antibody (mAb) cocktail: peridinin-chlorophyll protein Cy5 (PerCP-Cy5) conjugated lineage specific mAbs (eBioscience; San Diego, CA) (CD11c, NK1.1, CD11b, Ter119, CD3e, CD8a, CD19, TCRβ, TCRγ, B220, and Ger1), APC conjugated anti-Kit mAb (eBioscience; San Diego, CA), Phycoerythrin/Cy7 (PE/Cy7) conjugated anti-Sca-1 mAb (eBioscience; San Diego, CA), PE conjugated anti-Flt3 mAb (eBioscience; San Diego, CA). Dead cells (propidium iodide–positive) were excluded from the analysis. HSCs were defined as lineage negative, Sca-1 and Kit positive, Flt3 negative cell population.

### Staining of paraffin sections

Five-micron paraffin spleen sections were stained with a-GATA1 Ab, clone D52H6 (Cell Signaling) and counterstained with hematoxylin. Antigen was retrieved at low pH.

### Identification of hematopoietic lineages in bone marrow chimeras

Single-cell suspensions were prepared from bone marrow from age- and gender-matched GF and SPF mice. Recipient SPF BALB/cJ mice irradiated at 900 rad received 2 × 10^6^ BM cells intravenously. Nine days after the transfer, the mice were sacrificed. Their spleens and bone marrow were fixed in Bouin’s fixative, sectioned, and stained with hematoxylin and eosin.

### RNA sequencing

RNAseq analysis was performed on splenic RNA from infected (preleukemic, sore 1) and uninfected GF and SPF BALB/cJ males and females (4 groups of mice, 6-8 mice per group). Splenic RNA was subjected to Illumina Next Gen Sequencing to generate a Directional Total RNA library. Ribosomal RNA was removed using a Ribo-Zero rRNA removal kit (Epicentre; Madison, WI). Gene expression was quantified through the Kallisto software (Bray et al., 2016). To identify genes regulated by both the microbiota and the virus, we performed a series of heuristic gene set filtering operations (see explanation in the text) using log-fold change >0.5 and *p* < 0.05 Wilcoxon rank sum test. Gene filtering and statistical analysis was conducted using MATLAB (R2019b) (Natick, MA).

### Measurement of serum IL-6

Serum IL-6 levels were assessed in uninfected, pre-leukemic, and leukemic mice using a Cytometric Bead Array for mouse Th1, Th2, and Th17 cytokines (BD Biosciences; San Jose, CA) as per manufacturer’s instructions.

### RNA isolation from spleens and quantitative reverse transcription PCR (RT-qPCR)

RNA was isolated from spleens using either guanidine thiocyanate extraction and CsCl gradient centrifugation (Chirgwin et al., 1979) or Purelink RNA Mini Kit (Invitrogen). Complementary DNA (cDNA) was generated using SuperScript IV Reverse Transcriptase (Life Technologies). Real-time PCR was conducted either with TaqMan Master Mix (ThermoFisher) or SYBR green master mix (Bio-Rad; Hercules, CA). TaqMan primer pairs and probes to amplify VSig4 (Mm00625349_m1), Serpinb9b (Mm00488405_m1), Rnf128 (Mm00480990_m1), and Beta-Actin (Mm02619580_g1) were from Applied Biosystems. The following forward and reverse primers were used to amplify VSig4 - 5’ GGAGATCTCATCAGGCTTGC3’ and 5’CCAGGTCCCTGTCACACTCT; Rnf128 - 5’TAGCTGTGCTGTGTGCATTG3’ and 5’GAATGTCACACTTGCACATGG3’; Serpinb9b - 5’AGCAGACCGCAGTCCAGATA3’ and 5’GTCTGGCTTGTTCAGCTTCC3’, and beta-actin - 5’GTATCCTGACCCTGAAGTACC3’ and 5’TGAAGGTCTCAAACATGATCTG3’ in SYBR green master mix reactions. cDNA was quantified using the comparative Ct method.

### ELISA

An enzyme-linked immunosorbent assay (ELISA) was used to detect anti-MuLV antibodies in MuLV-infected GF and SPF BALB/cJ mice as previously described (Case et al., 2008). Briefly, RL-MuLV virions isolated from infected SC-1 cells were treated with 0.1% Triton X-100 and bound to plastic in borate-buffered saline overnight. Two percent ovalbumin was used as blocking component. Sera samples were incubated at 4°C for one hour at a 1 x 10^-2^ dilution, followed by goat anti-mouse IgGs coupled to horseradish peroxidase (HRP) (SouthernBiotech, Birmingham, AL). Background optical density (OD450) values from incubation with secondary antibody alone were subtracted from values acquired from sera of infected mice.

## Results

### Microbiota is not required for MuLV transmission or replication

We have previously shown that the replication of a different retrovirus (Mouse Mammary Tumor Virus, MMTV) was dependent on the presence of the host microbiota, which served as a source of lipopolysaccharide (LPS) used by MMTV to dampen the host’s immune response. If a similar scenario (control of replication by the microbiota) was applicable to MuLV, it would be difficult to study a potential role of the microbiota in development of MuLV-induced pathology. To address the issue, we used BALB/cJ mice either treated with microbiota-depleting doses of antibiotics (Abx) (Kane *et al*., 2011) or reared in germ-free (GF) conditions. SPF and GF mice were infected by intraperitoneal (i.p.) injection of 0.22μM filter-sterilized MuLV as adults (G0 mice) and bred to produce offspring (G1 mice). SPF G1 mice, which were weaned either on regular or Abx-containing water, were examined for the presence of infectious virus along with GF G1 mice. This experimental design allowed us to test whether the microbiota is required for both virus replication and transmission. Frequency of infected cells and viremia were compared *via* infectious center and plaque assays, respectively (Rowe *et al*., 1970). Although Abx had no effect on viral transmission in SPF mice (Figure 1A), there was potential of an outgrowth of an Abx-resistant bacteria during the course of the experiment. However, GF MuLV-infected mice also showed similar frequency of infected cells in the spleens (Figure 1A) and infectious virions in the plasma (Figure S1A) compared to infected SPF mice. Thus, the system was robust for testing the role of the microbiota in leukemogenesis caused by MuLV.

**Figure 1:**
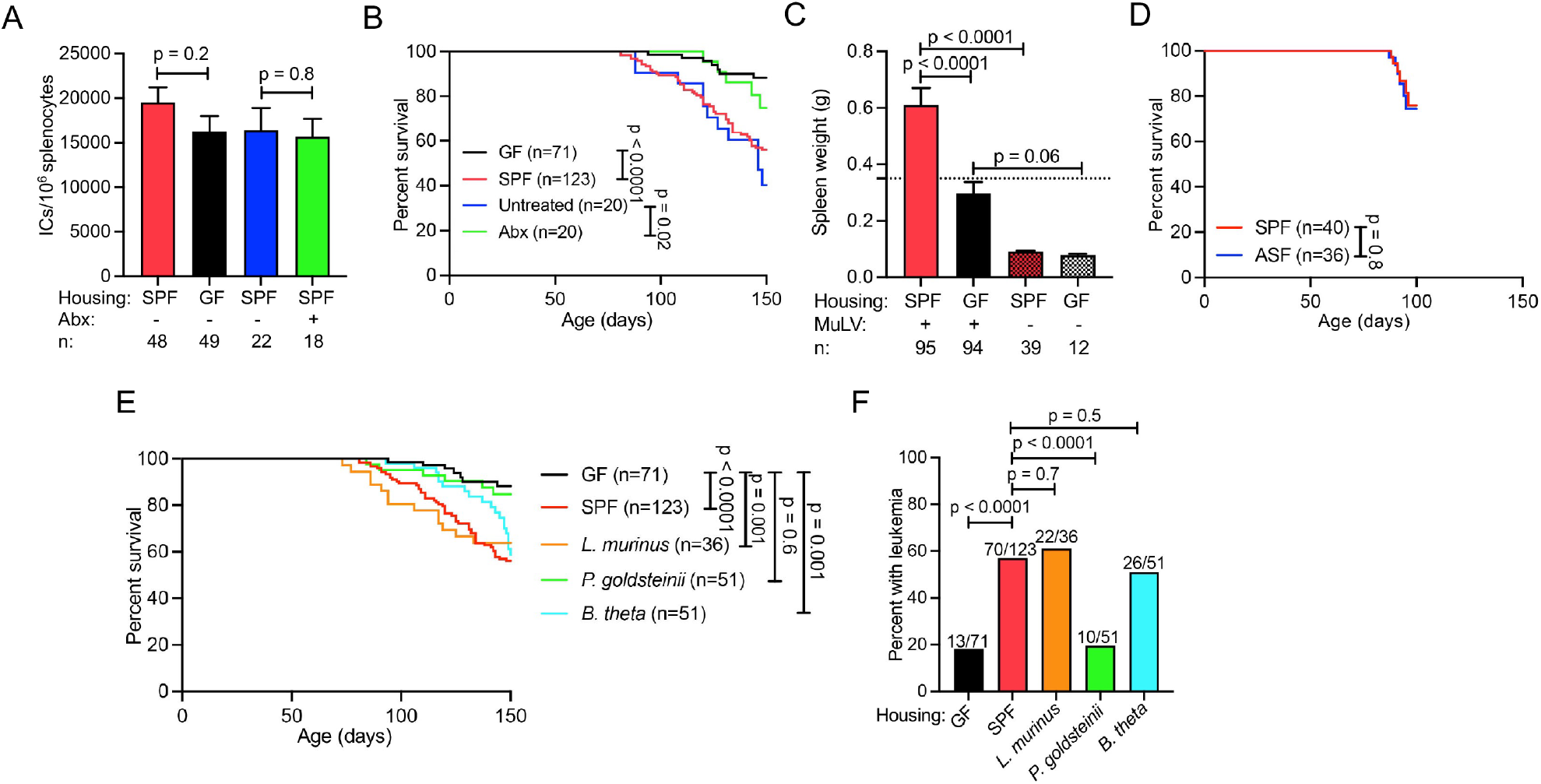
Dependence of MuLV-induced leukemia development on commensal bacteria. Adult BALB/cJ females from different experimental groups were injected with MuLV and bred (G0 mice). Their offspring (G1 mice) were monitored for leukemia. Diseased mice removed from the cohorts and mice surviving up to 150 days were examined according to a leukemia scoring system based on histological analysis of the spleen (Figure S1C). **(A)** Comparison of viral load [number of infectious centers (ICs) per 10^6^ splenocytes] in SPF, GF, and Abx-treated animals. IC assay was done on preleukemic mice with score 1. **(B)** Survival curves of G1 mice from the same groups. **(C)** Spleen weights of infected and uninfected SPF and GF mice at 4-5 months of age. Dotted horizontal line indicates 0.35g. **(D)** Survival curves of MuLV-infected SPF and ASF-colonized BALB/cJ mice observed for 97 days. **(E)** Survival of GF, SPF and gnotobiotic BALB/cJ mice colonized with single bacterial lineages monitored for 150 days. **(F)** Final assessment of leukemia development in mice from these groups at day 150. Abx, antibiotic. n, number of mice used per group. *p* values were calculated using an unpaired *t* test **(A, C),** Fisher’s exact test **(F)** and Mantel-Cox test **(B, D, E)**. Error bars indicate standard error of the mean.

### Microbiota accelerates virally-induced leukemia

Upon RL-MuLV infection, hematopoiesis within the bone marrow is blocked, resulting in compensatory extramedullary hematopoiesis (EMH) in the spleen (Hook *et al*., 2002). EMH leads to expansion of target cells susceptible for infection. RL-MuLV readily replicates within the rapidly proliferating erythroid progenitor cells, increasing the likelihood of proviral integration near a cellular proto-oncogene, a step necessary for development of erythroid leukemia as this virus does not encode for an oncogene (Figure S1B).

To test whether the microbiota has an effect on RL-MuLV pathogenesis, mice from different groups were infected by i.p. injection of filter-sterilized virus as adults (G0 mice) and bred to produce infected offspring (G1 mice) that were further observed for leukemia development. G1 mice, which received a physiologically relevant infectious virus dose from their parents, were aged and monitored for leukemia. Diseased mice removed from the cohorts and mice surviving up to 150 days were examined according to a leukemia scoring system which was based on histological analysis of the spleen. Uninfected mice retained splenic architecture and were given a score of 0. Pre-leukemic mice were defined as having increased extramedullary hematopoiesis and were given a score of 1. Leukemic mice exhibiting regions containing leukemic blasts with high mitotic activity were scored as 2, and more advanced cases, where the splenic architecture was wholly disrupted by immature leukemic blasts, were scored as 3 (Figure S1C). Leukemia score directly correlated with the spleen weight and all animals with the spleen weight greater than 0.35g had leukemia (score 3) (Figure S1D).

Infected SPF BALB/cJ mice exposed to water containing a broad-spectrum Abx cocktail since weaning as well as infected GF BALB/cJ mice had increased latency of the disease and markedly reduced spleen weights compared to infected untreated SPF mice (Figures 1B, 1C, and S1E). As leukemia resistant phenotype of Abx-treated and GF mice was not due to a reduction in viral replication (Figures 1A and S1A), it became obvious that the microbiota significantly augmented viral pathogenesis.

### Gut commensal bacteria differ in leukemia promoting properties

The commensal microbiota is composed of various microorganisms including bacteria, viruses, archaea, fungi, and unicellular eukaryotes. To define a subset of the microbiota capable of enhancing leukemia susceptibility of SPF mice, GF mice were colonized with a defined consortium of mouse commensal bacteria, Altered Schaedler’s Flora (ASF) (Dewhirst et al., 1999), bred and infected with filter-sterilized virus (G0 mice). Virus fate and pathogenesis was followed in G1 mice generated from infected G0 females. ASF restored high susceptibility of GF mice to leukemia (Figures 1D and S1F), proving that commensal bacteria promote virally-induced leukemogenesis.

ASF contains seven bacterial species that include both gram-positive and gramnegative lineages (Sarma-Rupavtarm *et al*., 2004). A single gram-positive bacterium, *Lactobacillus murinus* (*L. murinus*), and a single gram-negative bacterium, *Parabacteroides goldsteinii* (*P. goldsteinii*), were isolated from ASF and used to colonize GF mice. Progeny from *L. murinus* colonized females developed leukemia at a similar rate and incidence as infected SPF mice (Figures 1E and 1F). Conversely, progeny from infected *P. goldsteinii* colonized females exhibited leukemia development similar to that of infected GF mice (Figures 1E and 1F). Importantly, infected SPF and *P. goldsteinii* colonized mice had similar frequency of infected splenocytes (Figure S1G). Non-member of ASF, gram-negative commensal *Bacteroides thetaiotaomicron (B. theta)* also conferred leukemia susceptibility to GF infected mice (Figures 1E and 1F), although with a slight delay in disease development. ‘These data provide evidence that leukemia promoting factors are not inherent to all bacteria but are unique to certain bacteria across different bacterial phyla.

### Erythroid differentiation is stalled at a specific stage of development in infected SPF mice

To study the promotion of leukemogenesis by commensal bacteria, we sought to determine the stage of erythroid leukemia development that is halted in infected GF mice. Extramedullary hematopoiesis (EMH) is a result of MuLV-induced proliferation of hematopoietic stem cells (HSCs) in the spleen and occurs in 100% of RL-MuLV-infected BALB/cJ SPF mice (Hook *et al*., 2002). Both infected SPF and GF mice exhibited a robust expansion of HSCs, identified as the Sca-1^+^ population within nucleated spleen cells (Figure 2A), indicating that this phase in disease development was not affected in GF mice. In addition, BALB/cJ GF mice did not have defect in frequency of HSCs within the bone marrow (BM) (Figure 2B) or in differentiation of HSCs into hematopoietic lineages upon transfer into lethally irradiated mice (Figure 2C and Figure S2).

**Figure 2:**
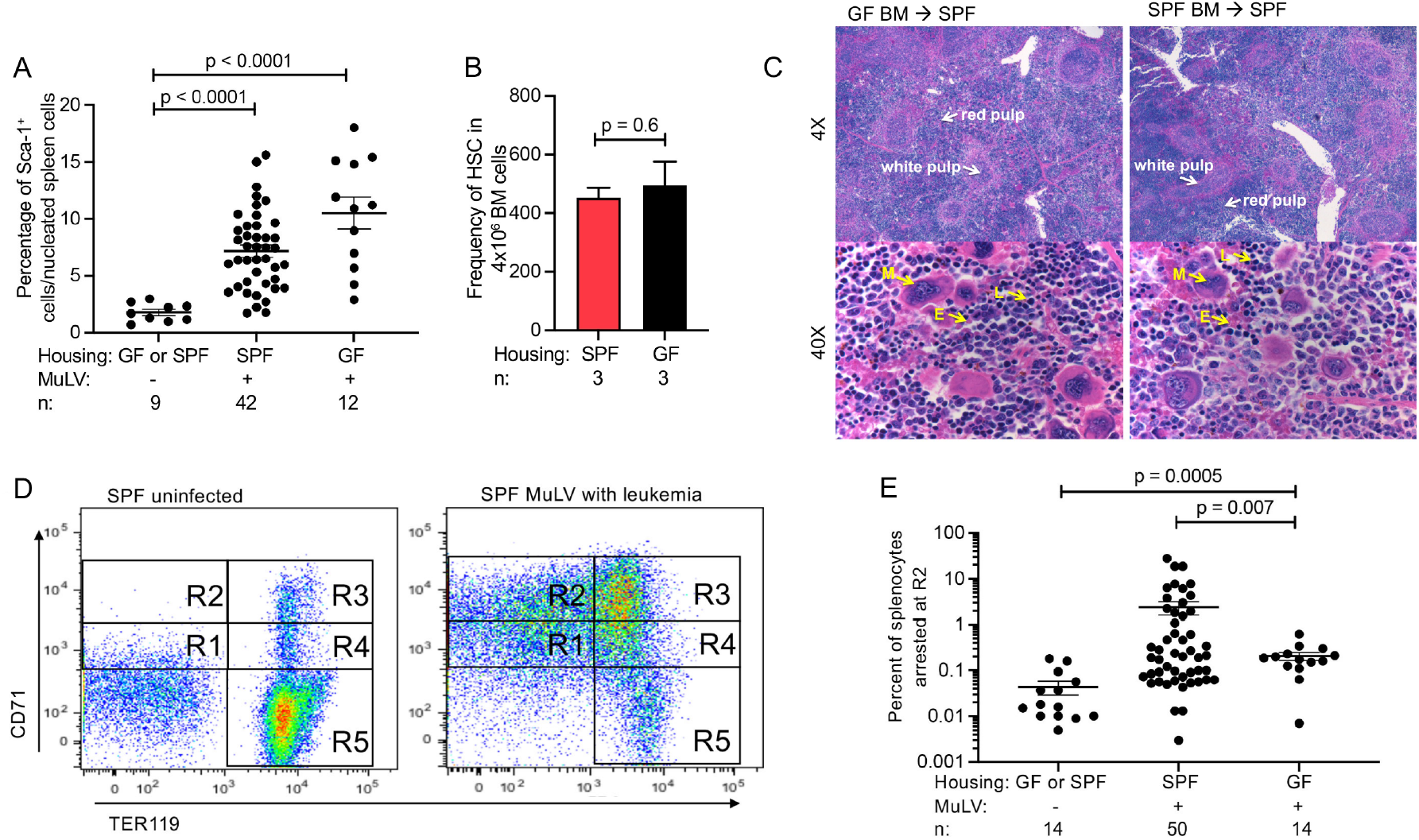
Comparison of leukemia development in SPF and GF BALB/c mice. **(A)** Increase in extramedullary hematopoiesis in the spleens of infected mice. Nucleated splenocytes were stained with anti-mouse Sca-1 mAb and analyzed by FACS. Uninfected GF and SPF BALB/cJ mice were used as negative controls. **(B)** Enumeration of HSCs (defined as lineage negative, Sca-1^+^ and Kit^+^, Flt3^-^ cell population) in the bone marrow of SPF (red bar) and GF (black bar) BALB/cJ mice. **(C)** Comparison of hematopoietic cells in the spleens of irradiated SPF BALB/cJ mice reconstituted with bone marrow transplants from GF and SPF BALB/cJ donors. Hematoxylin/eosin staining. Red and white pulp indicated by white arrows. Cells of the different hematopoietic lineages are identified by yellow arrows. Representative images of one of three recipient mice per group. M: megakaryocyte, E: erythroid precursor, L: lymphocyte. **(D)** Determination of a specific developmental stage at which erythroid cells are stalled in infected SPF mice. The stages of erythrocyte maturation were determined as described (Zhang *et al*., 2003) by staining nucleated spleen cells with a mixture of anti-mouse CD71/anti-mouse Ter119 mAbs. Staining profiles of indicated mice. **(E)** A quantitative comparison of cell numbers arrested at R2 (see **D**) stage in SPF and GF mice. n, number of mice used. *p* values calculated using unpaired *t* test. Error bars indicate standard error of the mean.

To further define the stage where disease development is blocked in GF mice, erythrocyte maturation in infected and uninfected SPF and GF mice was assessed by FACS analysis. During erythrocyte development, precursor cells initially express high levels of the transferrin receptor, CD71. As erythroid cells mature, they increase expression of erythrocyte-specific marker (Ter119) and subsequently decrease CD71 expression (Zhang *et al*., 2003). Five distinct populations of cells in Figure 2D are defined by their characteristic staining patterns: CD71 ^med^TER119^low^ (gate R1), CD71 ^high^TER119^low^ (gate R2), CD71^high^TER119^high^ (gate R3), CD71^med^TER119^high^ (gate R4), and CD71^low^TER119^high^ (gate R5) during erythroid maturation (Zhang *et al*., 2003). Splenocytes from MuLV-infected SPF mice were stalled at stage R2 of developmental process (Figures 2D and 2E). In contrast, significantly fewer infected GF mice have their spleen cells stalled at stage R2 (Figure 2D and 2E). This illustrates that the development of erythroid precursors is halted in in the presence of the virus and microbiota.

### Proinflammatory cytokine IL-6 is dispensable for leukemia development

Tumor promotion is often linked to chronic inflammation, which is defined as a prolonged, aberrant protective response to a loss of tissue homeostasis (Medzhitov, 2008). Inflammation-driven tumor promotion activates transcription factors in premalignant cells, which induce genes stimulating cell proliferation and survival (Grivennikov et al., 2010). One candidate factor that contributes to inflammation and tumorigenesis, is the proinflammatory cytokine IL-6. IL-6 has been shown to be overexpressed by host cells within the tumor microenvironment and by tumors of various etiologies (Lippitz, 2013), and also shown to be microbially induced and necessary for pre-leukemic myeloproliferation in genetically susceptible mice (Meisel et al., 2018). To determine the role of IL-6 in virally-induced leukemia, IL-6-sufficient and -deficient female mice were infected with RL-MuLV and their progeny were monitored for leukemia. IL-6-sufficient and - deficient mice both developed leukemia at the same latency and with similar incidence (Figure 3A). Interestingly, while about 50% of infected SPF BALB/cJ WT mice develop leukemia by 150 days (Figure 1B), we noticed that nearly 80% of SPF BALB/c IL-6^-/-^ mice exhibit leukemia by 150 days (Figure 3A). This could be attributed among other possibilities to the difference in strain background between BALB/cJ (the background strain used for all mice in the studies) and BALB/cByJ mice (the background strain of the IL-6^-/-^ mice) that could influence the microbiota altering the rate of leukemia development. We also measured levels of IL-6 in the sera of uninfected, infected, and leukemic mice only to find that the cytokine concentration was unchanged (Figure 3B). Therefore, in the case of RL-MuLV-driven leukemogenesis, the proinflammatory cytokine IL-6 can be ruled out as a contributing factor.

**Figure 3:**
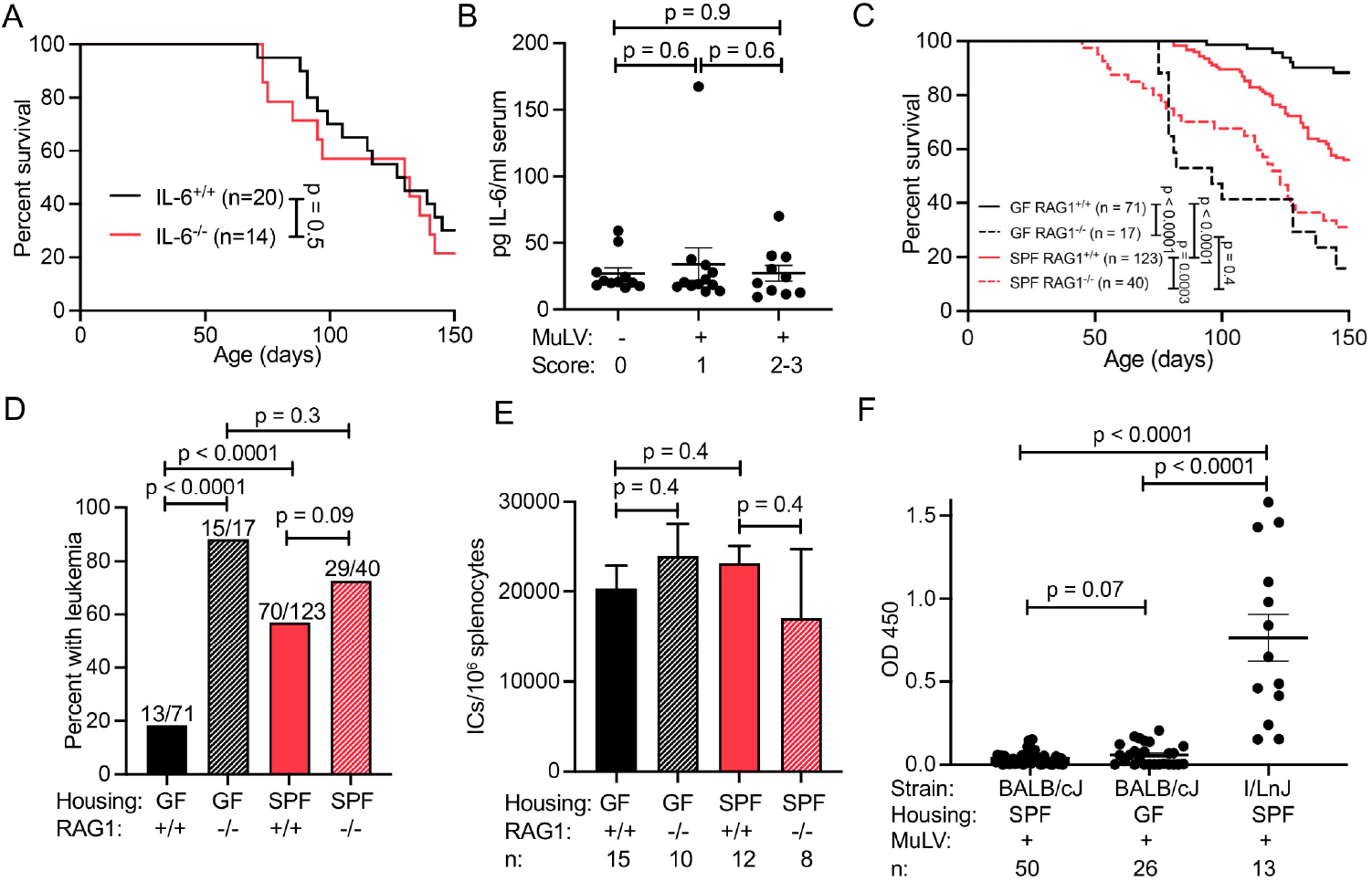
Comparison of leukemia susceptibility of immunosufficient and immunodeficient infected GF and SPF BALB/cJ mice. **(A)** Survival of IL-6 sufficient and IL-6 deficient MuLV-infected SPF BALB/cJ mice. **(B)** Serum cytokine concentration of IL-6 was measured in uninfected (score 0), infected preleukemic (score 1) and infected leukemic (scores 2-3) BALB/cJ mice using a flow cytometry bead-based assay. **(C)** Survival of infected RAG1-sufficient SPF, RAG1-deficient SPF, RAG1-sufficient GF, and RAG1-deficient GF BALB/cJ mice during 150 days. **(D)** Total leukemia incidence of infected indicated mice at 150 days. **(E)** Comparison of the viral burden (frequency of infected cells per 10^6^ splenocytes) in indicated preleukemic mice (score 1). **(F)** MuLV-specific ELISA to detect anti-virus antibodies. I/LnJ mice fostered on MuLV-infected BALB/cJ females are included as a positive control as they produce virus-neutralizing antibodies (Case *et al*., 2008). n, number of mice used. *p* values calculated using Mantel-Cox test **(A, C)**, Fisher’s exact test **(D)**, or unpaired *t* test **(B, E, F)**. Error bars indicate standard error of the mean.

### Commensal bacteria help tumors to escape the immune response

Maintenance of pre-cancerous and cancerous cells within the host requires the evasion or suppression of the immune response (Kim et al., 2007; Shankaran et al., 2001). To test the role of the adaptive immune response in resistance of GF mice to RL-MuLV-induced leukemia, we monitored leukemia development in immunodeficient RAG1 recombinase negative (lacking a functional adaptive immune system) GF BALB/cJ G1 mice born to infected RAG1-deficient G0 mice. RAG1 deficiency led to enhanced susceptibility of RL-MuLV infected GF mice to leukemia (Figures 3C and 3D). Importantly, both GF and SPF RAG1^-/-^ mice displayed a similar frequency of infected splenocytes, excluding the possibility that the sensitivity of RAG1-deficient mice to MuLV-induced tumors could be simply explained by an increase in viral replication in these mice (Figure 3E).

These data implied that increased leukemia susceptibility in SPF mice was a consequence of commensal bacteria suppressing the anti-tumor adaptive immune response. And, conversely, the absence of commensal bacteria in GF mice allowed an unsuppressed adaptive immune response to counteract the development of leukemia. Even though MuLV induces immunosuppression in SPF mice (Dittmer et al., 2004; Robertson et al., 2006), the immune response still controls leukemia development to a certain degree as SPF RAG1^-/-^ mice exhibited increased leukemia susceptibility compared with SPF RAG1-sufficient mice (Figures 3C and 3D). This response was unrelated to virus-specific antibodies (Abs) as neither infected GF nor SPF mice mounted virus-specific Ab responses (Figure 3F). In addition, significant susceptibility of GF RAG1^-/-^ mice to RL-MuLV-induced leukemia ruled out the possibility that resistance of GF wild-type mice to leukemia is due to the lack of provirus integration in the vicinity of oncogenes.

### MuLV infection and the microbiota produce a unique gene expression signature. To MuLV infection and the microbiota produce a unique gene expression signature

To identify microbiota-mediated signaling pathway(s) essential for counteracting the immune response controlling leukemia progression, RNAseq was performed on spleens from infected and uninfected age-matched GF and SPF mice (GF, GF MuLV, SPF, SPF MuLV) at the pre-leukemic stage (Figure 4A). Given the distinct phenotype of MuLV-infected SPF mice, we sought to identify gene expression changes associated with both microbiota and viral infection. A series of heuristic gene set filters were applied to the set of all genes across all mice. First, we assembled a set of genes that changed collectively between SPF MuLV mice and mice from the SPF, GF, and GF MuLV conditions (log-fold change > 0.5 and p 0.05 Wilcoxon rank sum test): intersection of (a) SPF MuLV vs SPF, (b) SPF MuLV vs GF, and (c) SPF MuLV vs GF MuLV (Figure 4A, positive gene set). Second, we assembled the group of genes from GF, GF MuLV, and SPF mice that differed from SPF MuLV condition (log-fold change > 0.5 and p 0.05 Wilcoxon rank sum test): union of (d) GF MuLV vs GF, (e) SPF vs GF MuLV, and (f) SPF vs GF (Figure 4A, negative gene set). Finally, we subtracted the negative gene set from the positive gene set, yielding candidate genes that were specifically influenced by the presence of both the virus and the microbiota. Knowing that leukemogenesis depends on the negative regulation of the immune response, we focused on genes that were known to have properties of negative immune regulation (Figure 4B, red arrows). Three genes - V-set immunoglobulin-domain-containing 4 (VSig4), serine (or cysteine) peptidase inhibitor, clade B, member 9b (Serpinb9b), and Ring finger protein 128 (Rnf128, also known as <u>g</u>ene <u>r</u>elated to <u>a</u>nergy in <u>l</u>ymphocytes, or GRAIL) were selected for further *in vivo* analysis.

**Figure 4:**
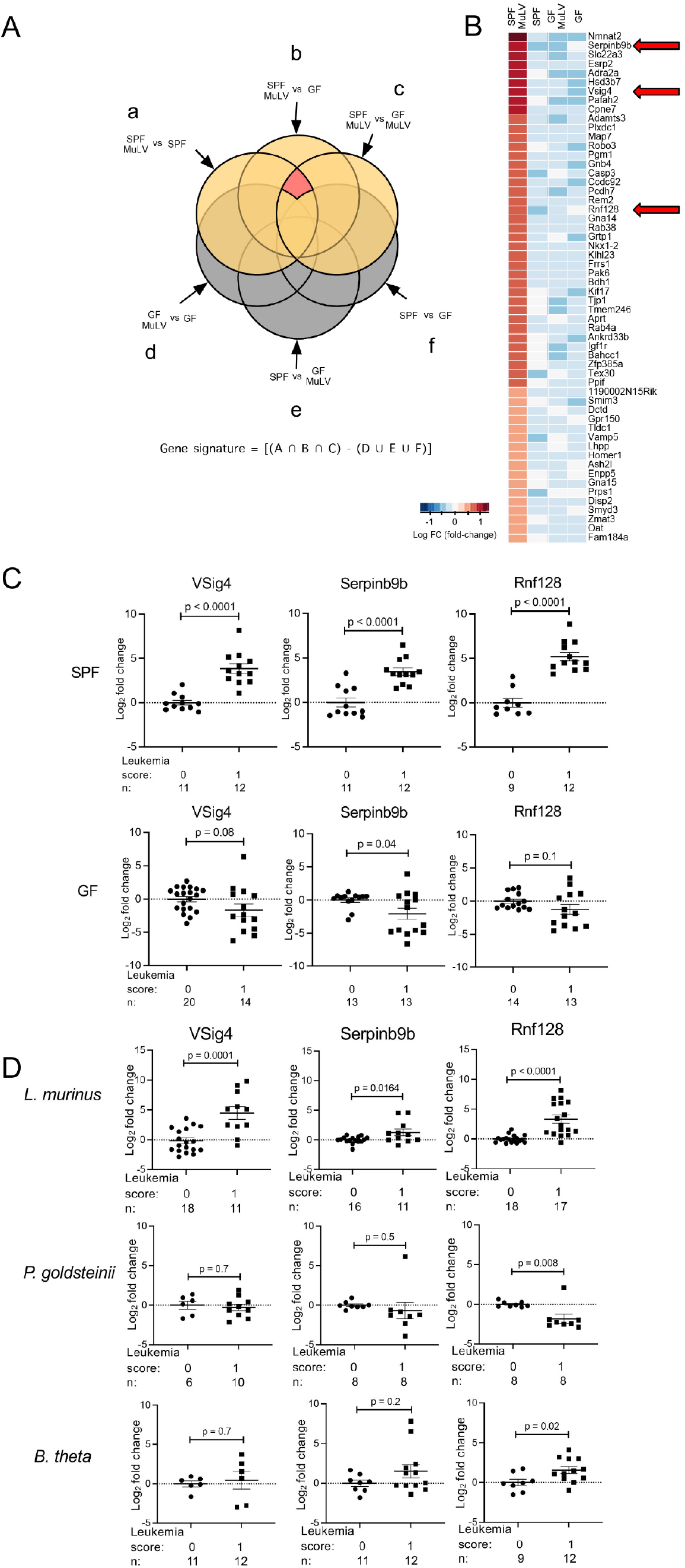
Induction of negative regulators of the immune response by commensal bacteria and the virus. RNA isolated from spleens of preleukemic mice (score 1) of 4 groups (SPF +/- MuLV infection and GF +/- MuLV infection) was subjected to high throughput sequencing. **(A)** Diagram detailing the series of operations taken to identify genes differentially expressed in spleens of mice from four groups. See explanation in the text. **(B)** Heat map of gene expression found to be significantly upregulated in SPF MuLV-infected mice compared to mice from all other groups. Red arrows indicate established negative regulators of adaptive immunity. **(C)** Real time quantitative PCR (qPCR) measurement of expression of the negative regulators of immune response in SPF and GF mice. Mice were either uninfected (score 0) or infected pre-leukemic (score 1). n, number of mice used. **(D)** qPCR with RNA isolated from spleens of uninfected (score 0) and infected pre-leukemic (score 1) mice colonized with *L. murinus, P. goldsteinii*, or *B. theta*. Data are represented as log2 fold change compared to uninfected controls, normalized to the endogenous control (beta-actin). Two-way ANOVA was used to identify genes upregulated in infected SPF mice compared to uninfected SPF, uninfected GF, and infected GF mice **(A, B)**. *p* values calculated using unpaired *t* test **(C, D)**. Error bars indicate standard error of the mean.

### Deficiency in Serpinb9b or Rnf128 is sufficient to confer leukemia resistance in SPF mice

VSig4 was discovered as a complement receptor and subsequently shown to inhibit T cells (Vogt et al., 2006; Zeng et al., 2016). Serpinb9b is a serine protease inhibitor that acts on and suppresses granzyme M (Bots et al., 2005). Rnf128 is a ubiquitin ligase that among many functions has been shown to ubiquitinate CD3 and CD40L on T cells, leading to their degradation (Lineberry et al., 2008; Nurieva et al., 2010). To confirm the upregulation of these genes by MuLV and the microbiota, real time quantitative PCR (RT-qPCR) was performed using RNA isolated from the spleens of pre-leukemic mice. Whereas SPF pre-leukemic mice showed significant upregulation of these genes compared to uninfected SPF mice (Figure 4C, top), GF pre-leukemic mice did not (Figure 4C, bottom). Notably, the expression of VSig4, Serpinb9b, and Rnf128 was also upregulated in the spleens of *L. murinus-*colonized mice but not *P. goldsteinii* colonized mice (Figure 4D) indicating that their induction correlated with the presence of bacteria with leukemia-promoting properties. Interestingly, Rnf128 was the only factor upregulated in the spleens of ex-GF mice colonized with leukemia-promoting *B. theta* (Figure 4D). Together, these data suggest that one or a combination of these negative immune regulators function in a microbiota-dependent fashion to promote leukemia development.

To provide a definitive proof that all or some of these genes (VSig4, Serpinb9b, and Rnf128) function to promote leukemia, a CRISPR-Cas9 approach was taken to generate BALB/cJ mice lacking these genes with the expectation that elimination of the critical factor(s) would result in leukemia resistance even in the presence of commensal bacteria. Guides were designed to target exon 1 of VSig4 (Figures S3A and S3D), exon 2 of Serpinb9b (Figure S3B), and the ring finger domain within exon 4 of Rnf128 (Figure S3C) resulting in frameshifting indels (VSig4 and Serpinb9b) and disruption of a functionally important domain of Rnf128 (Anandasabapathy et al., 2003). Mice with targeted mutations, their wild-type littermates and BALB/cJ mice bred in the same colony were injected with the virus and further bred to produce infected offspring monitored for leukemia. VSig4-deficient mice developed leukemia at a similar latency and incidence as VSig4-sufficient mice (Figures 5A–5B and Figure S3E), indicating VSig4 by itself does not contribute to MuLV-induced leukemia development. Interestingly, latency and incidence of leukemia development in Serpinb9b-deficient SPF mice and Rnf128-deficient SPF mice were both significantly delayed and reduced compared to wild type mice (Figures 5A–5B and Figure S3F). Thus, Serpinb9b-deficiency and Rnf128-deficiency significantly reduced leukemia susceptibility in SPF conditions without affecting viral replication (Figure S3G).

**Figure 5:**
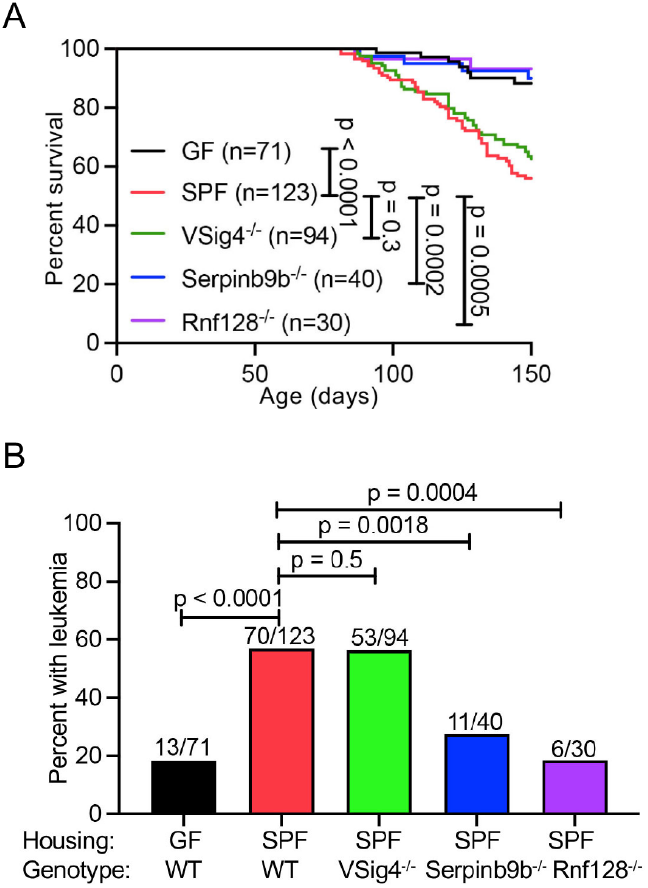
Resistance of Rnf128-deficient and Serpinb9b-deficient SPF mice to MuLV-induced leukemia. MuLV-infected BALB/cJ mice deficient in either VSig4, Serpinb9b, or Rnf128 were monitored for leukemia development. **(A)** Survival curves for up to 150 days are shown. **(B)** Final leukemia assessment at 150 days. n, number of mice used. *p* values calculated using Mantel-cox test **(A)** and Fisher’s exact test **(B)**. Error bars indicate standard error of the mean.

### Serpinb9b is upregulated via RIPK2-mediated signaling pathway

To ascertain how the host detects microbial derived factors which subsequently leads to upregulation of negative immune regulators, we investigated the role of host innate immune receptors in RL-MuLV-induced leukemia. Given that bacteria are sufficient to promote RL-MuLV induced leukemia, we investigated pattern recognition receptors that specifically detect bacterial moieties. We hypothesized that removal of the host sensor detecting leukemia-promoting bacteria would result in resistance to leukemia even in the presence of the microbiota. Mice lacking innate immune receptors TLR2, TLR4, and mice unable to signal through NOD1/2 via deficiency in adaptor RIPK2 were bred on the BALB/cJ background, infected with RL-MuLV and their offspring were monitored for leukemia. Mice deficient in TLR2, a receptor for bacterial products including lipoteichoic acid and capsular polysaccharide (de Oliviera Nascimento et al., 2012; Graveline et al., 2007; Han et al., 2003), displayed latency and incidence of leukemia similar to WT SPF mice (Figures 6A–6B). Similarly, TLR4-deficient mice, which are unable to detect LPS (Poltorak et al., 1998), exhibited leukemia latency and incidence similar to WT SPF mice (Figures 6A–6B). However, it was possible that signaling through either TLR2 or TLR4 could be redundant and promote leukemia development. To address this possibility, we infected TLR2^-/-^ TLR4^-/-^ mice and monitored their offspring for leukemia to find that they were as susceptible as mice deficient in either single receptor (Figures 6A–6B).

**Figure 6:**
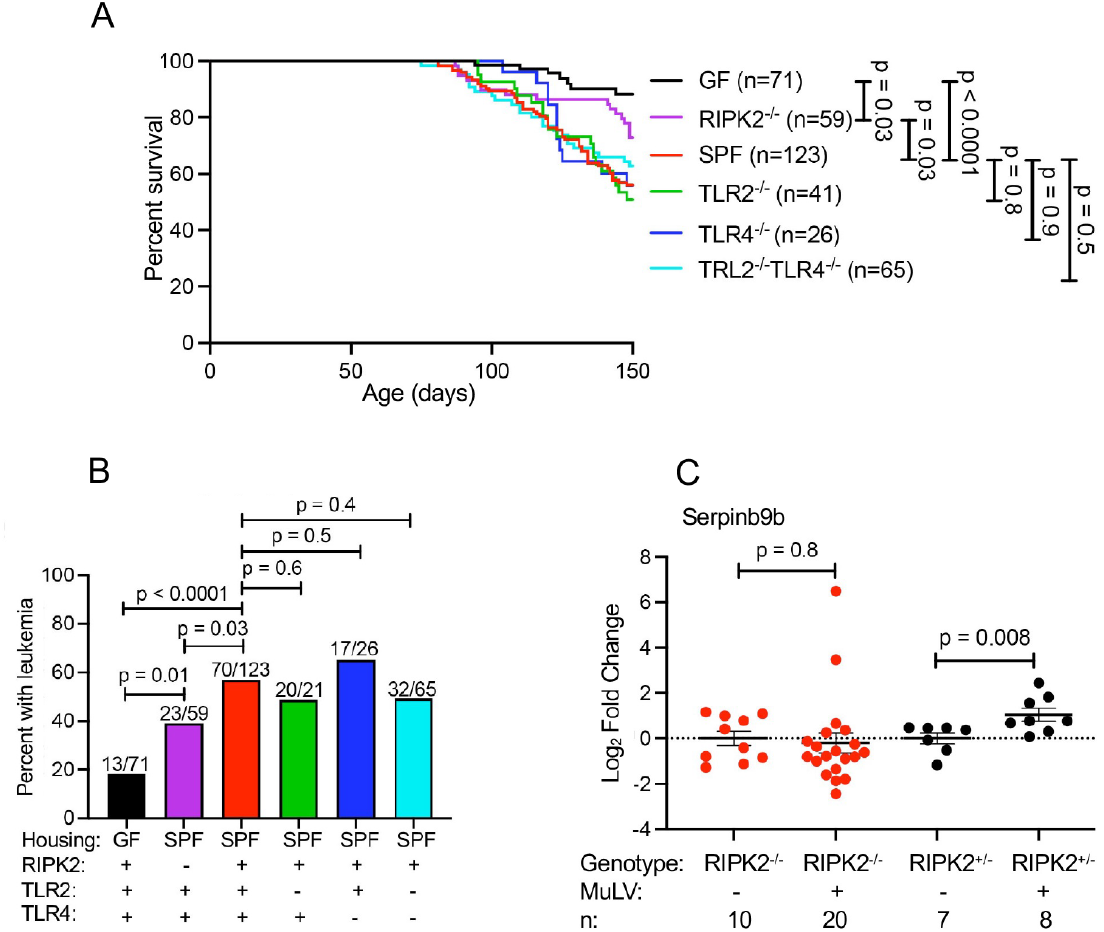
NOD1/2 adaptor RIPK2 contributes to MuLV-induced leukemia via upregulation of Serpinb9b. Leukemia development was monitored in RL-MuLV-infected SPF BALB/cJ mice deficient in either TLR2, TLR4, TLR2/TLR4, or RIPK2. **(A)** Survival curves up to 150 days. **(B)** Total leukemia incidence at 150 days. **(C)** Expression of Serpinb9b in splenic RNA from RIPK2^-/-^ and control RIPK2^-/-^ uninfected and infected mice analyzed by qPCR. Data are represented as log2 fold change compared to uninfected controls, normalized to the endogenous control (betaactin). n, number of mice used. *p* values calculated using Mantel-cox test **(A)**, Fisher’s exact test **(B)** or unpaired *t* test **(C)**.

At the same time, mice deficient in RIPK2, the downstream adaptor of intracellular peptidoglycan receptors NOD1 and NOD2 (Caruso et al., 2014), had significantly increased latency and decreased total leukemia incidence compared to WT SPF mice (Figures 6A–6B). These data suggested that detection of microbial products, likely intracellular peptidoglycan, and signaling through the NOD1/2 RIPK2 pathway promoted RL-MuLV induced leukemia development. Although RIPK2 deficient mice were significantly more resistant to leukemia development compared to WT SPF mice, RIPK2 deficiency did not confer the level of leukemia resistance as observed in WT GF mice (Figures 6A–6B). These data suggest that other innate immune sensors also play a role in RL-MuLV induced leukemia development.

Next, we tested which of the negative immune regulators upregulated in MuLV infected SPF mice were dependent on the RIPK2 signaling. Accordingly, RNA expression of the negative immune regulators in infected and uninfected RIPK2-deficient mice was compared. Whereas expression of Rnf128 and VSig4 was significantly upregulated in infected RIPK2-deficient mice (Figures S4A-S4B), expression of Serpinb9b was not (Figure 6C). Importantly, Serpinb9b was upregulated in infected RIPK2-sufficient control mice (Figure 6C). Together, these data demonstrate that activation of RIPK2 contributes to the upregulation of Serpinb9b in the presence of both, the virus and commensal bacteria. Therefore, a lack of Serpinb9b upregulation in RIPK2 deficient mice is likely the basis for the observed increase in leukemia resistance.

## Discussion

Retroviruses induce a broad range of tumors in vertebrates. In most cases retroviruses induce tumors by insertional activation of cellular proto-oncogenes during establishment of a provirus. Importantly, these proto-oncogenes are implicated in spontaneous tumors in humans. Although proto-oncogene up-regulation constitutes a necessary step for initiation of retrovirally-induced tumorigenesis, additional events - tumor promotion and progression (Kinzler and Vogelstein, 1996) - are required for successful tumor formation. Tumor promoters enhance cell survival and may induce clonal expansion of precancerous cells, whereas tumor progression is caused by additional mutations, which enable cancerous cells to invade neighboring tissues and metastasize.

The ability of the microbiota to regulate viral replication and pathogenesis caused by viruses belonging to different families (Baldridge et al., 2015; Kane *et al*., 2011; Kuss et al., 2011; Uchiyama et al., 2014) initiated our interest in its influence on MuLV-driven pathology. Some older literature suggested that the microbiota could influence MuLV-induced leukemia development, but it was highly controversial: germ-free animals infected with Friend’s MuLV (F-MuLV) either exhibited greater disease severity (Mirand and Grace, 1963) or milder disease (Isaak et al., 1988; Kouttab and Jutila, 1972) than their SPF control mice. Several factors may explain the discordance: at that time viral stocks were likely contaminated with Lactate Dehydrogenase Elevating Virus (LDV) (Dittmer et al., 2019), which is known to suppress and delay CD8^+^ T cell response against F-MuLV prolonging acute viral infection (Robertson et al., 2008). Furthermore, at that pre-PCR and sequencing era the sterility of the experimental isolators was tested only using culturing techniques (Kouttab and Jutila, 1972). Thus, unculturable bacteria could have been missed.

In our studies of retrovirally-induced leukemia that originates in an organ distant from the gut, we found that the intestinal commensal bacteria enhanced the leukemogenesis. In contrast to experiments in which microbiome supported viral replication (Baldridge *et al*., 2015; Kane *et al*., 2011; Kuss *et al*., 2011; Uchiyama *et al*., 2014), we found that the decreased leukemogenesis observed in RL-MuLV infected germ-free mice could not be explained by decrease in the virus burden. Instead, we discovered that the microbiota promotes leukemia development by negatively controlling the immune response. The immune system plays a role in controlling tumors of various etiology at each of the steps of tumor development (Vinay et al., 2015). Under its pressure, tumor cells exploit multiple mechanisms of avoidance of the immune responses: induction of regulatory T cells (Jacobs et al., 2012), defective antigen presentation due to down-modulation of antigen processing machinery (Hicklin et al., 1999; Johnsen et al., 1999), production of immunosuppressive mediators (Lind et al., 2004; Pasche, 2001) to name a few.

The high susceptibility and rapidity with which RL-MuLV infected GF RAG1^-/-^ mice develop leukemia supports a role for cells expressing somatically rearranged immune receptors in controlling tumor development in GF mice: T cells, B cells and cells carrying rearranged receptors but resembling innate cells, such as NKT cells. As neither SPF nor GF infected virus-susceptible mice produce neutralizing antibodies against viral antigens (Figure 3F), any prominent role for B cells in protective anti-tumor immune response seems unlikely. The absence of virus-neutralizing antibodies also explains the lack of control of extracellular virions in GF and SPF mice enabling the virus to spread from cell to cell. Therefore, the adaptive immune response against MuLV pathogenesis is likely controlled by T cells and/or NKT cells. Anti-tumor immunity response by NKT cells is thought to be primarily through their support of other effector cells like CD8^+^ T cells and NK cells via production of Th1 cytokine IFN-*γ* (Crowe et al., 2002) or IL-2 (Metelitsa et al., 2001). Both CD4^+^ T cells and CD8^+^ T cells play decisive roles in suppressing tumor development and growth (Toes et al., 1999). CD4^+^ T cells contribute to anti-tumor immunity by secretion of proinflammatory cytokines such as IL-2, IFN-*γ*, and TNF-*α* (Quezada et al., 2010). CD4^+^ T cells have also been reported to adopt cytotoxic activity by means of Fas-mediated cell death or perforin and granzymes induced apoptosis (Nagata and Golstein, 1995; Raskov et al., 2021; Xie et al., 2010). That said, RAG deficiency has been associated with reduced functionality of NK cells (Karo et al., 2014), suggesting that this cytotoxic cellular subset could also be targeted by microbiota-driven tumor promoting negative immunoregulation.

RNA-seq analysis revealed three negative immune regulators (VSig4, Rnf128, and Serpinb9b) which expression was significantly increased in the presence of both the virus and the microbiota. Rnf128 and Serpinb9b proved to be critical for leukemia development in RL-MuLV-infected SPF mice (Figure 5A). In line with our results, overexpression of the members of the ovalbumin family of serpins, to which Serpinb9b belongs, as well as Rnf128, are markers for poor prognosis in some human cancers [(Bai et al., 2020; ten Berge et al., 2002; Uhlen et al., 2017) and our own analysis of previously published data (Bolouri et al., 2018)]. A homolog for Serpinb9b has not been identified in humans, but related family member, human proteinase inhibitor 9 (SerpinB9) inhibits perforin-mediated cytotoxic T lymphocyte (CTL) cytotoxicity (Medema et al., 2001). Additionally, murine Serpinb9 was shown to protect mouse melanoma tumors from granzyme B mediated killing (Jiang et al., 2020). Published data support the idea that Serpinb9b functions in a similar manner by inhibiting the action of granzymes toward tumors (Bots *et al*., 2005). Rnf128 is highly expressed in anergic T cells and ubiquitinates key activation signaling molecules, such as CD3 and CD40L, resulting in their degradation (Lineberry *et al*., 2008; Nurieva *et al*., 2010). Rnf128 deficient primary CD4^+^ T cells do not develop anergic phenotype in various models and also exhibit hyperactivation upon TCR stimulation (Kriegel et al., 2009). Rnf128 deficiency in CD8^+^ T cells enhances their anti-tumor effector function by increasing expression of IFN-γ, granzyme B, perforin 1, and TNF-*α* (Haymaker et al., 2017). Precisely how Serpinb9b and Rnf128 function in the context of MuLV infection is yet to be discovered.

Although VSig4 is a *bona fide* negative regulator (Vogt *et al*., 2006), VSig4^-/-^ mice were as susceptible as wild-type littermates to RL-MuLV-induced leukemia (Figure 5A). It is possible that VSig4 serves as a marker of cells that have immunosuppressive function that is independent of VSig4. For example, VSig4^+^ resident macrophages have documented immunosuppressive function in various experimental models (Chen et al., 2011; Chen et al., 2010; Fu et al., 2012; Vogt *et al*., 2006; Xu et al., 2010).

Since bacterial products stimulate multiple innate pattern recognition receptors, it is important to understand which particular pathway(s) is important for promotion of leukemogenesis. Our studies illustrated a role for RIPK2, but not TLR2 or TLR4, in promoting MuLV induced leukemia development. In contrast to our studies, two prior reports, which examined the effect of NOD/RIPK2 pathway on tumors in the colon and liver demonstrated a suppressive effect of this signaling on tumor development (Chen et al., 2008; Ma et al., 2015). The mechanism by which activation of this pathway influenced tumor development was distinct in the two models of tumorigenesis. Erythroblastic leukemia originates from erythroid precursors found in infected spleens and the bone marrow (Hook *et al*., 2002), locations distant from the gut potentially explaining the discrepancies. RIPK2 likely supported leukemia development through upregulation of negative immune regulator Serpinb9b as its expression was increased in WT but not RIPK2-deficient MuLV-infected SPF splenocytes. RIPK2-Serpinb9b signaling axis does not entirely explain high susceptibility of SPF mice to leukemia compared to GF mice: the development of leukemia in RIPK2-deficient SPF mice was significantly reduced but did not reach the low level of GF animals (Figure 6B). Thus, additional signaling pathways must contribute to bacteria-dependent promotion of leukemogenesis. This pathway most likely involves Rnf128, which was the only negative regulator upregulated in infected *B. theta* colonized gnotobiotic mice susceptible to leukemia (Figure 4D).

Importantly, the bacterial factors promoting leukemia development are not intrinsic to gram-positive or gram-negative bacteria as members from both groups confer leukemia susceptibility. Furthermore, bacteria belonging to the same order of Bacteroidales, *P. goldsteinii* and *B. theta*, exhibit different leukemia-promoting capabilities. A major difference between the two Bacteroidales bacteria utilized in this study is the presence of an S-layer in *P. goldsteinii* (Fletcher et al., 2007). The S-layer is a bacterial cell structure composed of protein or glycoprotein thought to have evolved to provide protection against environmental factors (Fletcher *et al*., 2007; Sleytr et al., 1993) and the host’s immune response (Ezeji et al., 2021). Thus, S-layer may prevent proinflammatory signaling (Cuffaro et al., 2020; Kverka et al., 2011). and upregulation of the negative regulators in response to it. It seems obvious that commensals capable of downregulation of adaptive immune responses by upregulating the described negative regulators do so to maintain tolerance to themselves. Tumor promotion is obviously a side effect that requires additional genomic events stimulated by the retroviral infection.

In summary, our study highlights a role for the microbiota in the pathogenesis of a retrovirus in the absence of decreased virus replication. In this study, for the first time, we demonstrate that commensal bacteria facilitate leukemia development *via* induction of negative regulators of the immune response – a novel gut microbiota-mediated mechanism that enables tumor progression. As these negative regulators can be linked to a poor prognosis in certain human cancers (Bai *et al*., 2020; ten Berge *et al*., 2002; Uhlen *et al*., 2017), it is likely that similar mechanisms of immune evasion operates in tumors induced by viruses and potentially in spontaneous tumors of non-viral origin.

## Author contributions

J.S. performed most of the experiments reported in the paper. A.K. carried out computational analysis of the RNA-seq data. K.O. performed experiments with IL-6 and RAG1 deficient mice. J.W. conducted experiments with Abx treated mice, experiments on erythroid differentiation, and monitored ASF colonized mice. S.L. performed real time PCRs and IC assays for Serpinb9b^-/-^ and Rnf128^-/-^ mice. S.G. generated the leukemia scoring system, scored splenic tissue sections as well as sections of tumors that developed in Trp53-deficient and Wnt1 transgenic mice and contributed to experimental design. S.E. help to design CRISPR/Cas9 approaches to target Serpinb9b and Rnf128. A.J. cultivated *B. thetaiotaomicron*. M.F. and A.C. contributed to experimental design and the edit of the manuscript. J.S. and T.G. wrote the manuscript. T.G. conceived the project and performed many of the experiments.

## Acknowledgements

We thank members of the Golovkina and Chervonsky labs for helpful discussions, and Joseph Picard and Helen Beilinson for editing the manuscript, and Barbara Kee and Renee de Pooter for help with HSC staining. This work was supported by NIH grants, R21AI138224 to T.G. and by R01CA232882 to T.G and M.F. J.S. and J.W. were supported by T32 GM007183 and T32 AI007090, respectively. This work was also supported by NIH/NIDDK Digestive Disease Research Core Center grant DK42086 and NIH grant P30CA014599.

