## Supplemental Figures for "Gut commensal bacteria enhance pathogenesis of a tumorigenic murine retrovirus"

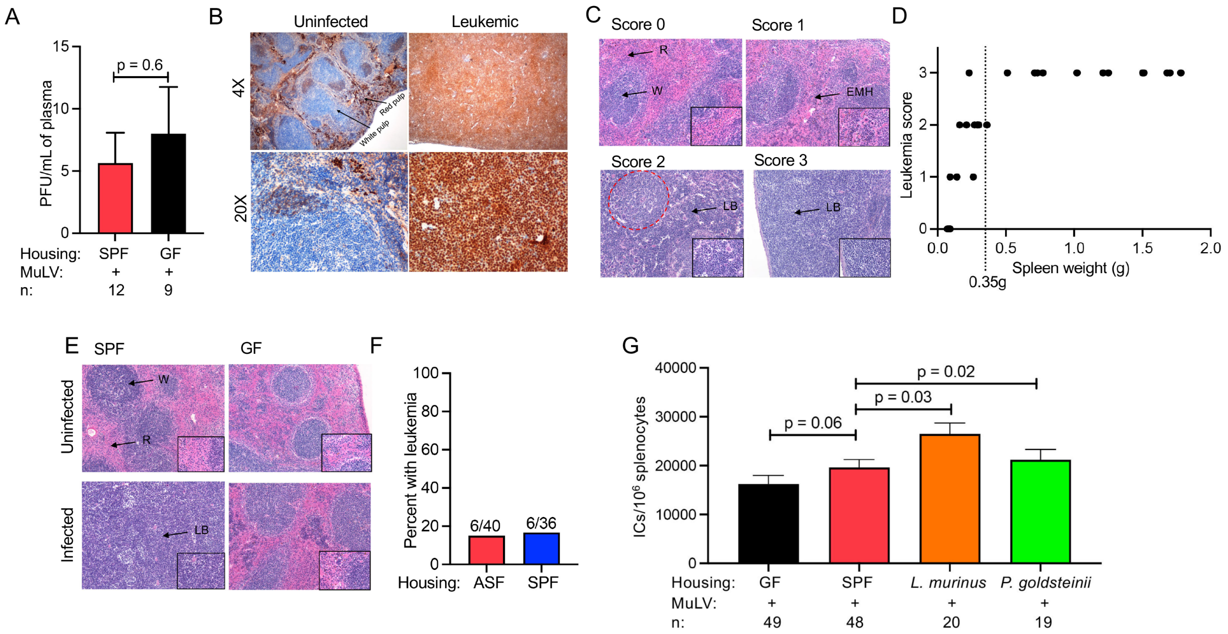


**Supplemental Figure 1: Infected GF mice exhibit minimal splenic tissue disruption, yet have similar viremia and splenic viral load compared to infected SPF and infected monocolonized mice. (A, B)** Female SPF BALB/cJ mice were injected with MuLV and their progeny were monitored for leukemia. **(A)** Plaque forming units (PFU) per mL of plasma. **(B)** Paraffin section of the spleen from an infected SPF mouse was stained with anti-GATA1 Ab and counterstained with hematoxylin. **(C)** Disease scoring system. Uninfected mice retain splenic architecture and were given a score of 0. R, red pulp; W, white pulp. Pre-leukemic mice were defined as having increased extramedullary hematopoiesis and were given a score of 1. Leukemic mice exhibiting regions containing leukemic blasts (LB) with high mitotic activity were scored as 2. Red dotted circle are the remnants of the white pulp. Score 3 was given to mice in which splenic architecture was wholly disrupted by immature LBs. Representative stained sections of spleens from SPF infected mice with indicated scores. Magnification - 10X, 40X magnification is shown in lower right corners. **(D)** Correlation of spleen weight with leukemia score determined by histological examination of hematoxylin and eosin (H & E) stained spleen sections. Vertical dotted line indicates spleen weight of 0.35g. **(E)** Example hematoxylin-and eosin stained-splenic sections from uninfected or infected SPF and GF mice at 4 months of age. Black arrows indicate red and white pulp. Magnification - 10X, 40X magnification is shown in lower right corners. **(F)** Total leukemic incidence in infected ASF-colonized and infected SPF mice monitored for 97 days. **(G)** Frequency of infected splenocytes (infectious centers, ICs) per $10^6$ splenocytes in SPF and gnotobiotic mice. n, number of mice used. *p* values were calculated using an unpaired *t* test **(A, G)** or Fisher's exact test (**F**). Error bars indicate standard error of the mean.

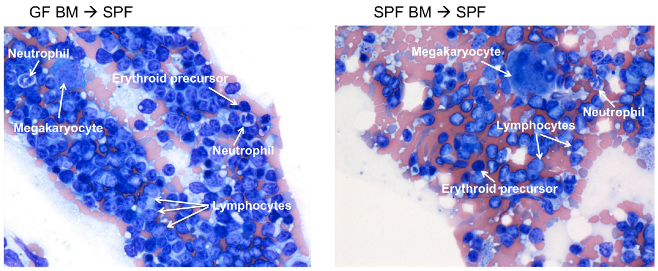


**Supplemental Figure 2: Hematopoietic stem cells from GF mice differentiate into all hematopoietic linages in the bone marrow of recipient mice.** SPF BALB/cJ mice were irradiated with 900 rads and subsequently received 2 x $10^6$ BM cells from either SPF or GF mice intravenously. Mice were euthanized nine days post transfer. Hematoxylin-and eosin stained smears of BM from irradiated SPF recipients of either GF (left) or SPF (right) BM.

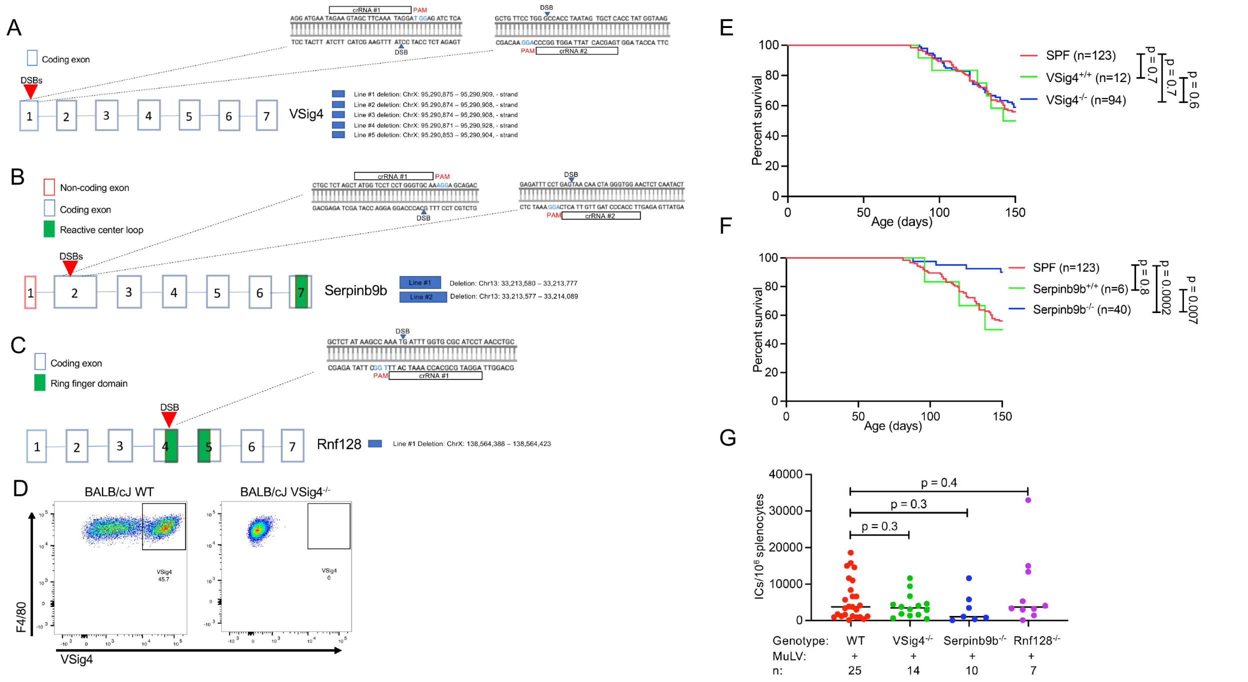


**Supplemental Figure 3: CRISPR-Cas9 targeting strategy for generation of VSig4-, Serpinb9b-, and Rnf128-deficient mice.** CRISPR-Cas9 targeting strategy for **(A)** VSig4, **(B)** Serpinb9b, and **(C)** Rnf128. Red arrow indicates location of double-stranded breaks within the targeted gene. Targeted sequences are shown on the right. All lines generated and the regions of the chromosome deleted are listed in blue boxes. **(D)** Peritoneal macrophages isolated from wild-type mice and VSig4$^{-/-}$ mice were stained with anti-VSig4 and anti-F4/80$^{+}$ antibodies. Representative FACS plot depicting staining in a wild-type mouse (plot on the left) and VSig4$^{-/-}$ mouse from line #2 (plot on the right). Deletion of VSig4 in the other lines was also confirmed by FACS (data not shown). **(E)** Survival curve of SPF MuLV-infected VSig4$^{-/-}$, VSig4$^{+/+}$ littermates, and wild-type non-littermate BALB/cJ mice bred in the colony. **(F)** Survival curve of SPF MuLV-infected SPF Serpinb9b$^{-/-}$, Serpinb9b$^{+/+}$ littermates, and wild-type non-littermate BALB/cJ mice bred in the colony. As Rnf128 is mapped to X chromosome we had limited Rnf128+/+ mice as littermate controls and thus, used non-littermate BALB/cJ mice bred in the colony as experimental controls. **(G)** Infectious centers per $10^6$ splenocytes from pre-leukemic SPF WT,

VSig4$^{-/-}$, Serpinb9b$^{-/-}$, and Rnf128$^{-/-}$ mice. n, number of mice used. *p* values calculated using unpaired *t* test **(G)** or Mantel-Cox test **(E, F)**. Error bars indicate standard error of the mean.

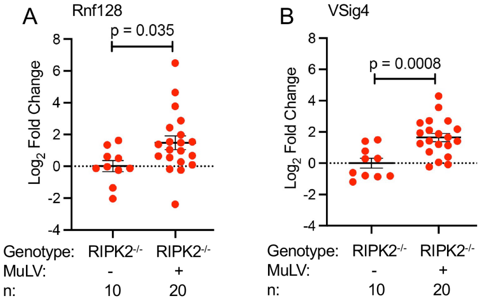


**Supplemental Figure 4: Upregulation of Rnf128 and VSig4 in infected SPF mice is independent of the NOD/RIPK2 pathway.** Expression of Rnf128 and VSig4 in splenic RNA from uninfected and infected RIPK2$^{-/-}$ mice analyzed by qPCR. Data are represented as $\log_2$ fold change compared to uninfected controls, normalized to the endogenous control (beta-actin). n, number of mice used. *p* values calculated using unpaired *t* test.
